# PiggyBac transposon and transposase delivered in *trans*, in *cis*, and as mRNA for engineering CAR-T cells: a multi-objective optimization

**DOI:** 10.64898/2026.10.16.758153

**Authors:** Nattanicha Deedit, Koramit Suppipat, Supannikar Tawinwung, Yozo Nakazawa, Nattiya Hirankarn, Thanyavi Chinsuwan

## Abstract

Chimeric antigen receptor (CAR)-T cell therapy has become a revolutionary strategy in cancer immunotherapy. However, conventional manufacturing strategies, which rely on viral vectors to deliver genetic material to host genomes, are hindered by many obstacles that have limited their widespread clinical application. This study aims to optimize the delivery of piggyBac transposon for simple and cost-effective manufacturing of CAR-T cells. We tested three different piggyBac delivery strategies: (i) conventional two-plasmid system (CONV), (ii) helper and donor one-plasmid system (ONE), and (iii) mRNA transposase in combination with a DNA donor plasmid (MRNA). Each method was evaluated for transfection efficiency, cell viability, and transgene copy number. The results demonstrated that, without enrichment, ONE approach generated CAR-T cells with the highest gene delivery efficiency at the cost of lower viability and higher transgene copy number, while transfection using MRNA offered a reduction in transgene copy numbers at the cost of lower efficiency. Interestingly, with enrichment by autologous feeder, the shortcoming in viability was completely overcome while retaining higher efficiency and higher transgene copy number. The result of this study lays the groundwork for a more efficient and safer piggyBac-engineered CAR T cell manufacturing process, and it should facilitate the decision in method selection for further optimization and clinical scale-up.

**Significance statement:** This study compared the efficiency, viability, and transgene copy number of CAR-T cells made using piggyBac transposon system delivered in three different formats: in *trans* as two plasmids (CONV), in *cis* as one plasmid (ONE), and transposase as an mRNA plus a transposon plasmid (MRNA). All three formats are being used in different clinical trials worldwide. While studies comparing ONE against CONV and MRNA against CONV exist and each demonstrates better results for their format against CONV, no three-way comparison study exists.

## Introduction

Chimeric antigen receptor (CAR)-T cell therapy has had great success in hematologic malignancies, such as diffuse large B cell lymphoma (DLBCL) and relapsed/refractory B cell acute lymphoblastic leukemia (B-ALL).^1,2^ As of 2026, US Food and Drug Administration (FDA) and European Medicines Agency (EMA) have approved seven CAR-T therapies for the treatment of hematologic cancers.^3,4^

Conventionally, CAR-T cells are transduced using viral systems, but there are also non-viral platforms, including transposon systems, that can stably integrate transgenes into the host genome. Non-viral systems have been developed for CAR-T manufacturing to overcome issues associated with viral systems such as high cost of viral vectors, limited insert size, complex regulatory requirements, and potential immunogenicity.^5^ In contrast, transposon systems are less costly, less complex, and are able to accommodate a much larger cargo size.^5,6^ When manufactured without pre-activation, piggyBac transposon-engineered CAR-T cells also contain a sizeable proportion of cells with memory phenotypes.^7–9^ PiggyBac transposon was first isolated from the cabbage looper moth *Trichoplusia ni*.^10^ The conventional method of transgene delivery using piggyBac transposon is to deliver the components *in trans*: the donor plasmid carries the transgene of interest while the helper plasmid encodes the transposase gene. Since then, more delivery formats have been developed to improve the transfection efficiency. To avoid the need for helper plasmids, Urschitz *et al.* have proposed delivery *in cis*, whereby the transposase and the transposon are delivered in the same plasmid.^11^ The strategy increased the probability that a cell would receive both the transposase and the gene of interest, ultimately improving transfection efficiency. In another report, Kaštánková *et al.* have proposed delivering the transposase as *in vitro*-transcribed (IVT) mRNA combined with the CAR transgene as linear DNA.^12^ The researchers showed the transfection efficiency to be higher than that of the conventional two-plasmid system. While both formats were demonstrated to be better than the conventional method, no three-way comparison has been made.

In this project, we compared the transfection efficiency, cell viability, and transgene copy number (TCN) of three different piggyBac delivery systems: (i) helper-and-donor one-plasmid system (“ONE”), (ii) IVT mRNA piggyBac transposase combined with one donor plasmid (“MRNA”), and (iii) conventional two-plasmid system (“CONV”), using FMC63-based anti-CD19 CAR-T as a model.

## Results

### Different formats exhibited significantly different efficiency-viability-TCN trade-offs at equivalent doses

To examine the inherent characteristics of different delivery formats (Figure 1A), we first manufactured the CAR-T cells without feeder cell enrichment (Figure 1B). T cells were isolated, activated, and transfected at various nucleic acid concentrations. They were expanded for 7 days to ensure sufficient dilution of the plasmids and that the expression would mostly derive from the integrated transgene.

**Figure 1:**
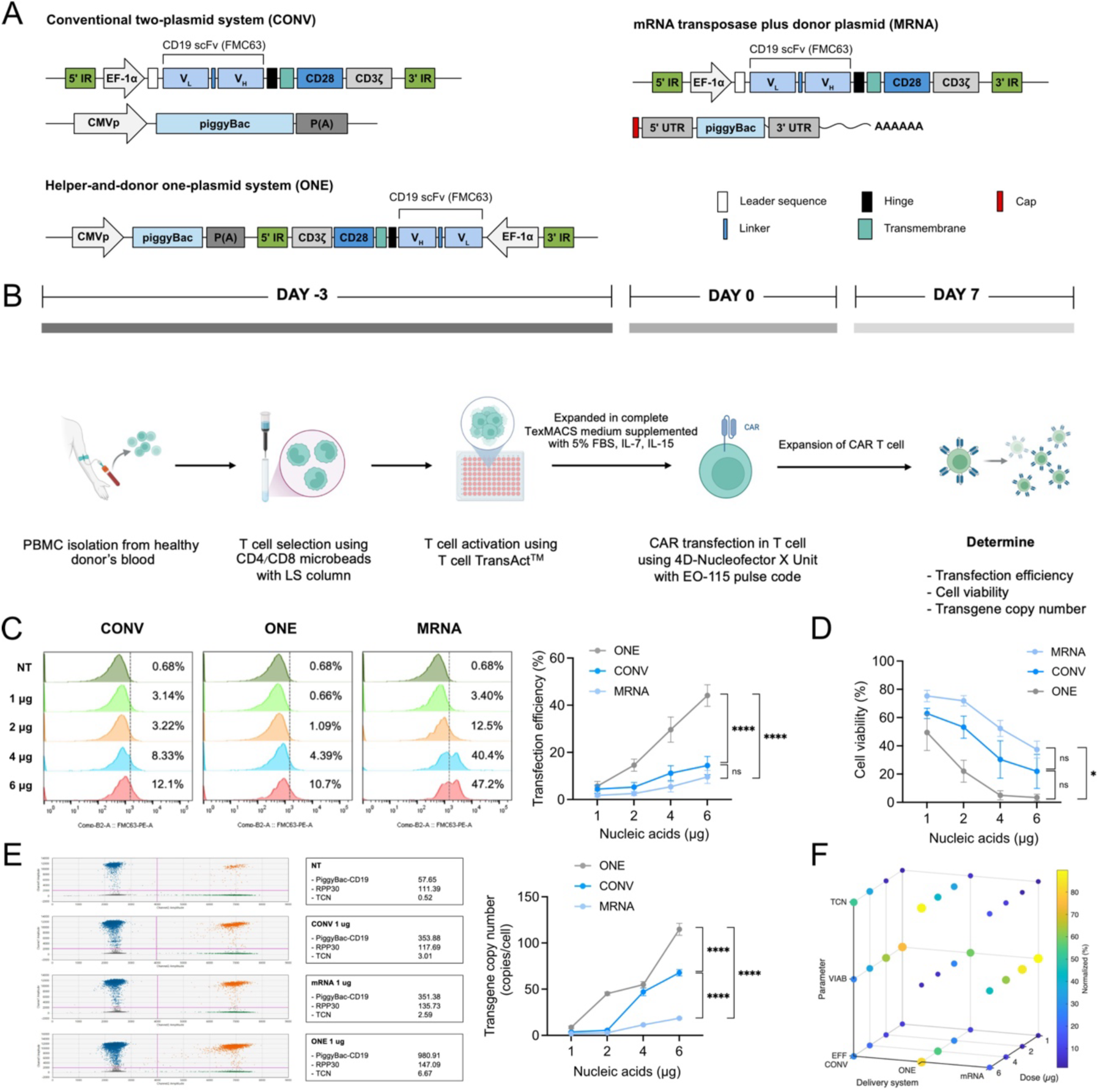
Efficiencies, viabilities, and transgene copy numbers achieved with different delivery formats of piggyBac in CAR-T cells produced without enrichment by feeder. (A) Schematic of piggyBac delivery formats. (B) Experimental design for engineering piggyBac CD19 CAR-T cells under feeder-free condition. (C–E) The parameters concerned as measured9 on day 7 post-transfection, showing representative (left) and summary (right) of transfection efficiencies (C), cell viabilities (D), and representative (left) and summary (right) of transgene copy numbers (E). (F) Three-dimensional bubble plot showing normalized average values of the three parameters across four doses of the three delivery formats. Data are presented as mean ± standard error of the mean (SEM), n=4. Data in C–E were analyzed by two-way ANOVA followed by Tukey’s multiple-comparisons test. ns (not significant), *p<0.05, **p<0.01, ***p<0.001, ****p<0.0001.

All delivery formats exhibited positively dose-dependent increases in transfection efficiency. Notably, ONE format showed the highest transfection efficiency compared to the others (Figure 1C). Conversely, all formats also caused negatively dose-dependent decreases in cell viability, with the ONE format causing significantly more cell deaths than other approaches at the equivalent doses. On the other hand, MRNA format retained a high level of cell viability in every nucleic acid concentration tested (Figure 1D). The read-out on TCN mirrored that of efficiency: ONE format led to more copies being integrated into the genome, positively correlating with the dose supplied (Figure 1E). CONV format settled in between the two in terms of efficiency-viability-TCN trade-off.

In summary, ONE format showed the greatest transfection efficiency but ranked lowest in TCN and viability parameters compared to the other formats. On the contrary, mRNA transposase plus one donor plasmid showed a better safety profile, but had low transfection efficiency (Figure 1F).

### Expressions of exhaustion markers were influenced by the delivery formats

We also evaluated exhaustion marker expression across the three approaches. For PD-1, TIGIT, and LAG-3, significantly higher expression was seen in ONE format at the higher doses (Figure 2A). TIM-3 expression did not differ between the three approaches at any concentration.

**Figure 2:**
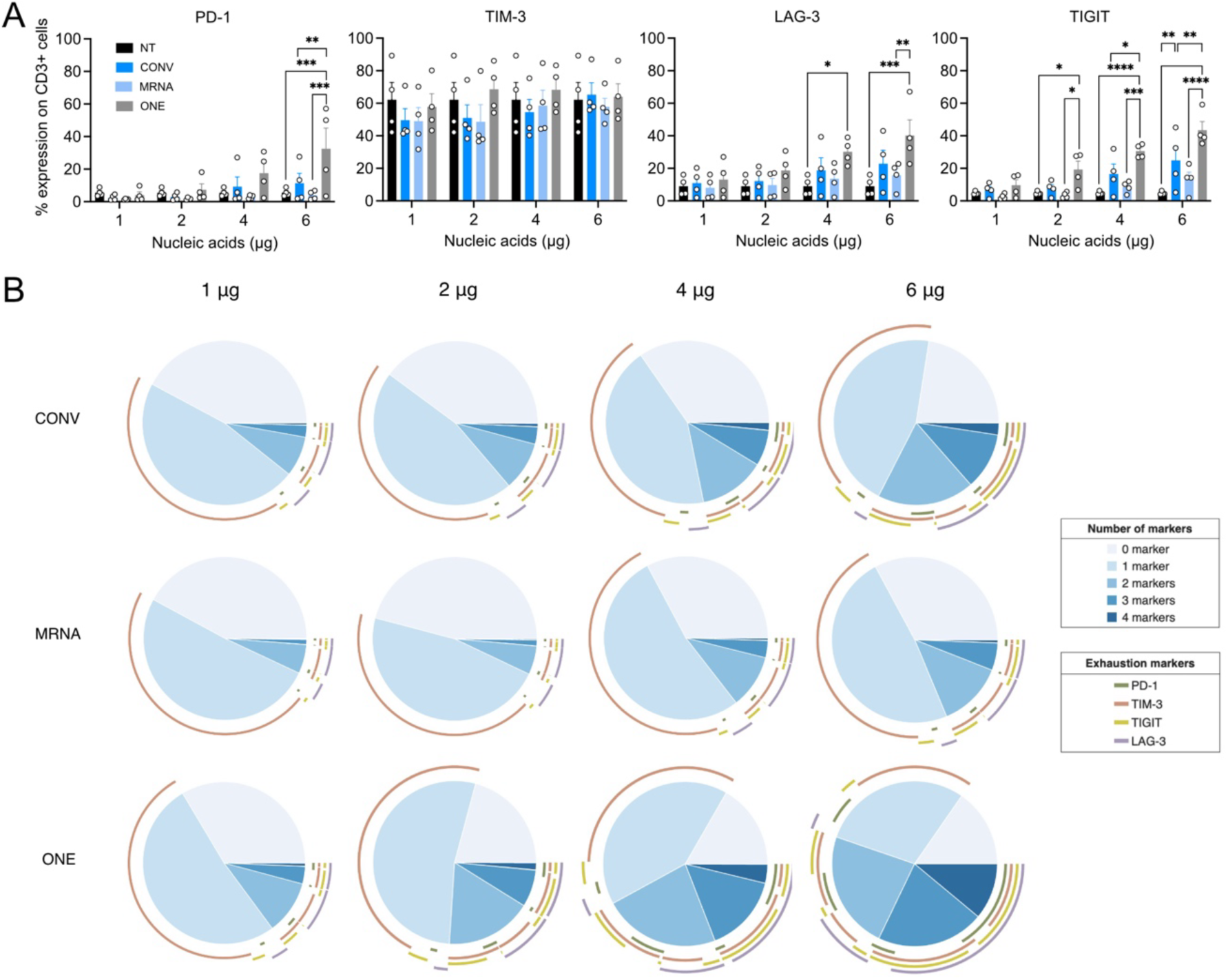
Expression of selected exhaustion markers on the engineered CAR-T cells (A) Expression of PD-1, TIM-3, LAG-3, and TIGIT on day 7 post-transfection on CD19 CAR T cell made by different delivery (B) Pie chart representing co-expressions of exhaustion markers. Data are presented as mean ± standard error of the mean (SEM), n=4. Data were analyzed by Two-way ANOVA followed by Tukey’s multiple-comparisons test. ns (not significant), *p<0.05, **p<0.01, ***p<0.001, ****p<0.0001.

Since co-expression of several exhaustion markers signifies worse functionality of T cells,^13^ we examined the proportions of CAR-T cells concomitantly exhibiting 0–4 of the markers above. The ONE format again displayed the lowest fraction of cells with 0 exhaustion markers across all four doses. This fraction fell from 33.55% at 1 µg to 15.35% at 6 µg of nucleic acid, and the combined 2–4 marker co-expression fraction was the largest among all formats (Figure 2B).

Unlike the exhaustion phenotype, neither the delivery formats nor the dose of nucleic acids altered the proportions of memory phenotype (Figure S1A) or CD4/CD8 (Figure S1B).

Altogether, it can be concluded that high concentrations of nucleic acid affect exhaustion marker expression, and the ONE format increases this risk more than the other approaches.

### The correlation between nucleic acid concentrations and the three parameters

To further analyze the gains and losses of the three parameters achieved by varying the nucleic acid concentration, linear regressions were fitted to the data. We found that the positive linear correlation between nucleic acid dose and transfection efficiency had the steepest slope in ONE approach with a relatively good fit (R^2^ = 0.83), while the remaining two were similarly and vastly shallower (Figure 3A). Notably, the inverse correlations between nucleic acid concentration and cell viability showed similar slopes but with different Y-intercepts (Figure 3B). For TCN, the analysis revealed a clear delineation, where the ONE format gained TCN heavily as the dose increased, while that of the MRNA format was much shallower (Figure 3C).

**Figure 3:**
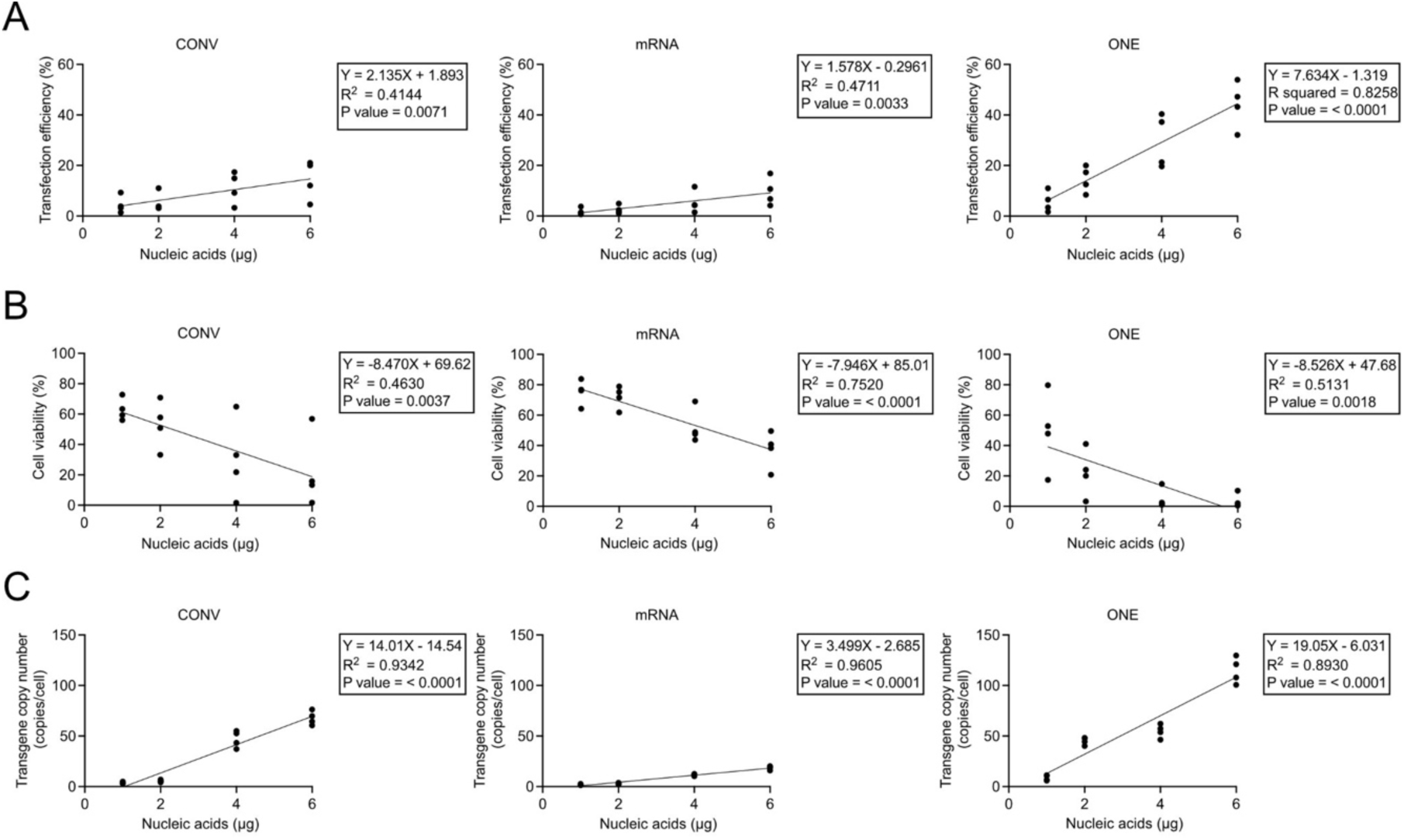
Correlation between nucleic acid concentration and the three parameters under feeder-free condition. (A) Correlation between nucleic acid concentration and transfection efficiency. (B) Correlation between nucleic acid concentration and cell viability. (C) Correlation between nucleic acid concentration and transgene copy number.

On closer examination, the slopes of nucleic acid against efficiency and against TCN tracked similarly for ONE and MRNA, but not for CONV. In other words, as more nucleic acid was supplied, ONE and MRNA gained 2.2–2.5 transgene copies per 1 percent increase in efficiency, whereas CONV gained 6.6.

Taken together, these analyses show that both MRNA and ONE can be interpreted as better than CONV depending on the prioritized parameters, supporting earlier studies.^11,12^ MRNA approach appears safer with gentler titration, while ONE approach maximizes efficiency but needs careful titration to avoid genotoxicity and cell loss.

### Identifying the optimal trade-off using ideal point analysis and the minimum Euclidean distance

Ideal point and minimum Euclidean distance were used to determine the optimal nucleic acid concentration for each approach. The ideal point for that respective format was defined as the within-group maxima for efficiency and viability and the within-group minimum for TCN. The Euclidean distance was then calculated using the average over all donors. The doses with the lowest Euclidean distances were 2 μg, 2 μg, and 1 μg for CONV, MRNA, and ONE approaches, respectively (Figure 4A).

**Figure 4:**
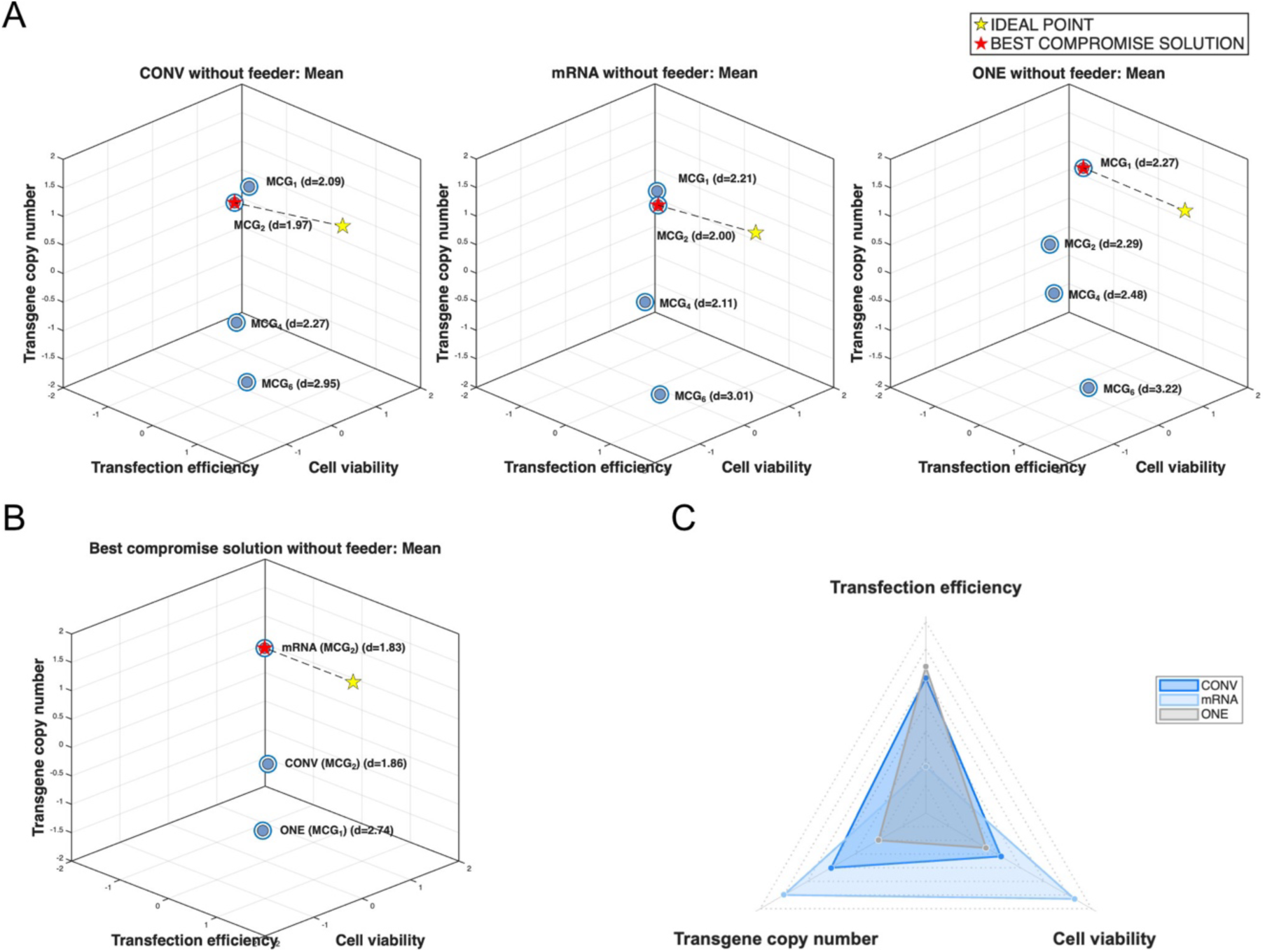
Ideal point and Euclidean distance analysis of the optimal trade-offs between the three parameters exhibited by the three delivery formats. (A) Ideal point and Euclidean distance analysis for optimal nucleic acid concentration of each delivery system. (B) Ideal point and Euclidean distance analysis for optimal delivery system. (C) Radar chart representing the performance of each delivery method at its optimal concentration for three parameters.

Once the optimal concentration was established for each of the delivery systems, another round of comparison was performed. In this round, the overall extrema were used to define the ideal point instead. Following the method described, the best compromise solution under the feeder-free condition was achieved using MRNA format (Figure 4B). The performance of each delivery approach at its optimal concentration was summarized in Figure 4C.

### Enrichment with feeder cells resolved the viability shortcoming of the ONE format

After establishing the optimal dose for each delivery format, those doses were used to investigate the performance of each approach using irradiated autologous feeder cells, a more practical manufacturing protocol (Figure 5A).^8,14,15^

**Figure 5:**
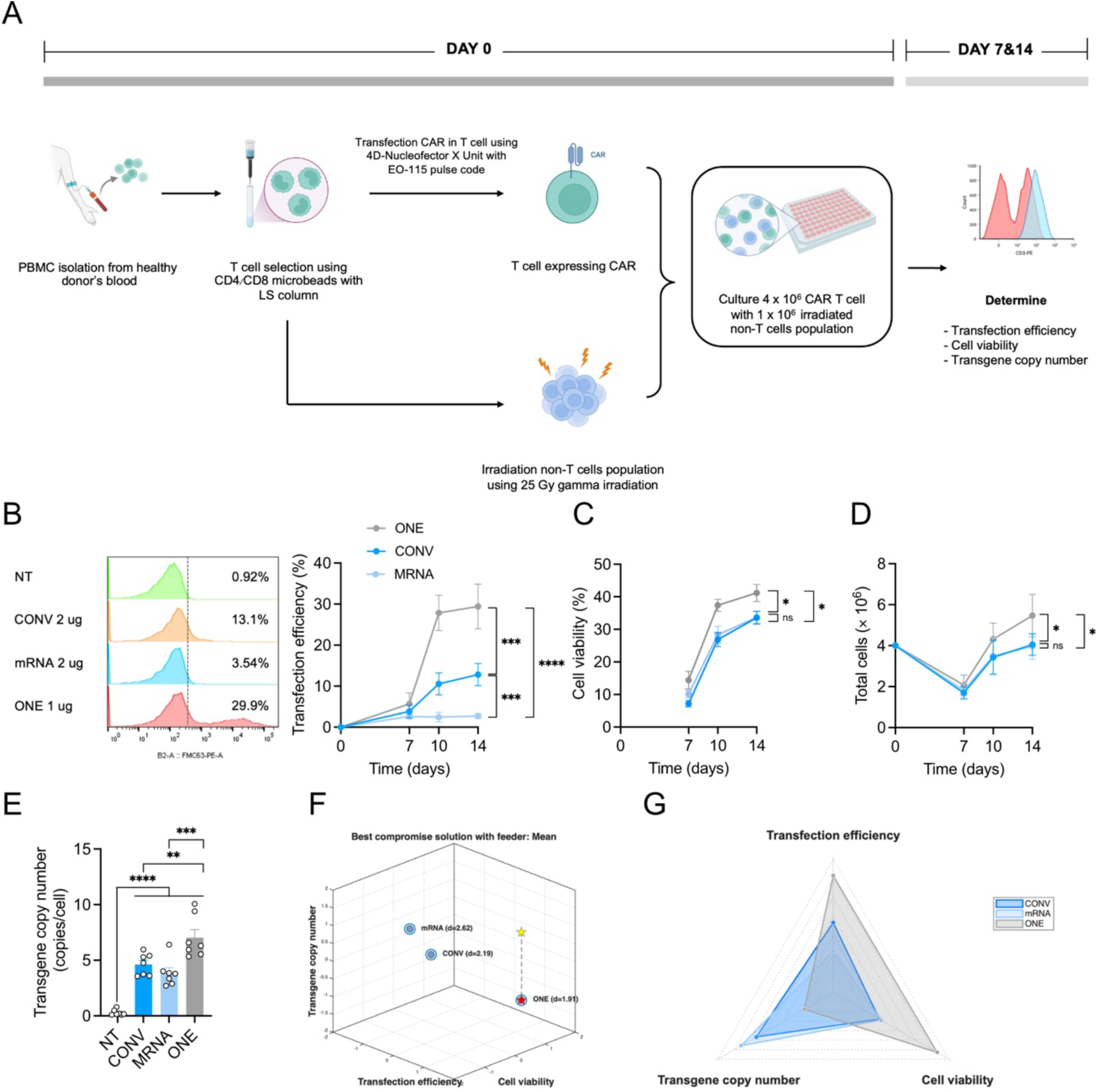
Efficiencies, viabilities, cell expansions, and transgene copy numbers achieved with different delivery formats of piggyBac when the CAR-T cells were enriched with feeder cells. (A) Experimental design for engineering piggyBac CD19 CAR-T cells with feeder enrichment. (B– D) On days 7, 10, and 14, the cells were assayed for transfection efficiency (B), cell viability (C), and total cell count (D). (E) Transgene copy number was measured at end point on day 14. (F) Ideal point and Euclidean distance analysis for optimal delivery system. (G) Radar chart representing the performance of each delivery method at its optimal concentration when three parameters were weighed together. Data are presented as mean ± standard error of the mean (SEM), n=7. Data in (B), (C), (D), and (E) were analyzed by two-way ANOVA followed by Tukey’s multiple-comparisons test. ns (not significant), *p<0.05, **p<0.01, ***p<0.001, ****p<0.0001.

We found that the transfection efficiency gradually increased from day 7 to 14 as the cells recovered after electroporation. On day 14, the transfection efficiencies for CONV, MRNA, and ONE were 12.8%, 2.71%, and 30.9%, respectively (Figure 5B). Likewise, cell viability also recovered from day 7 to 14 (Figure 5C). Surprisingly, the cell viability of the ONE format was the highest, which was corroborated by the total cell count (Figure 5D). On the contrary, there was no difference between CONV and MRNA regarding viability and cell expansion. However, the inherent property of ONE format regarding TCN persisted even under feeder enrichment, and it was still higher than the other two (Figure 5E and Figure S2).

The reversal of viability consequently changed the result of the ideal point analysis. In this round, ONE format had a considerably shorter Euclidean distance to the ideal point when all three attributes were considered across the group average. This suggests that with feeder enrichment, ONE approach represents the best compromise solution compared to the other approaches (Figure 5F). The performance of each delivery system at optimal concentration among the three parameters was summarized in Figure 5G.

In a similar manner to the feeder-free experiment, the phenotypes of piggyBac-engineered CAR-T cells across the three delivery systems were assessed. The results showed no differences between the three formats in TIM-3, TIGIT, or LAG-3 expression levels. However, the MRNA format exhibited a slightly higher PD-1 level (Figure S3A). In contrast to the feeder-free experiment, all three formats did not show different co-expression of various exhaustion markers (Figure S3B). The composition of memory phenotypes and the CD4/CD8 ratios did not differ either (Figure S3C and D).

In sum, these findings suggest that the expansion protocol can considerably alter the trajectory of cell recovery following electroporation during *ex vivo* expansion, as evidenced by the reversal of viability results for the ONE approach. Under a more practical expansion scheme, ONE approach emerged as the optimal compromise solution.

## Discussion

In the present study, three non-viral piggyBac transposon-engineered CAR T cell production was compared: (i) conventional two-plasmid system (CONV), (ii) a helper and donor one-plasmid system (ONE), and (iii) mRNA transposase plus one donor plasmid (MRNA). We demonstrated that CD19 CAR-T cells could be successfully generated from all three piggyBac delivery formats. To compare them, we assessed the capability and possible risk profile of each approach using three main parameters: transfection efficiency, cell viability, and transgene copy number. We also examined exhaustion markers, immunophenotype, and T cell subsets. Production was performed under feeder-free stimulation first to evaluate the intrinsic nature of the product attained by each delivery format. Later, the optimal dose of each format was chosen and used, with a larger sample size, for generating CAR-T cells with feeder enrichment, mimicking an actual protocol.

We found that CAR T cells stimulated with CD3 and CD28 agonists under the feeder-free condition altered the immunophenotype: these CAR T cells had a higher T_EMRA_ fraction, which indicates late differentiation or poor quality and is associated with poor expansion. In contrast, CAR T cells cultured with feeder cells had a higher T_CM_ fraction and a lower T_EMRA_ fraction, indicating that feeder cells support a more favorable immunophenotype.^15^

Under both conditions, ONE delivery provided the highest transfection efficiency across nucleic acid concentrations. Transfection efficiency was dose-dependent in all three formats. With feeder cells, efficiency improved for ONE and CONV. In particular, ONE provided the highest transfection efficiency and *ex vivo* expansion when produced with feeder enrichment, making it well suited to high-yield CAR T cell production. However, it also produced the highest transgene copy number, suggesting that further optimization is warranted if clinical production is the goal. In terms of potential risk of malignant transformation, transposase delivered as an mRNA has been shown to be safer than the other systems.^12,16^ This observation is reflected in our study by the consistently low TCN achieved by MRNA delivery, with or without enrichment.

Cell viability decreased with increasing nucleic acid concentration for all three delivery systems, which indicates that higher nucleic acid doses are more toxic to the cells. Cell death was highest in ONE, particularly at higher doses, whereas MRNA had the highest viability across all doses. This suggests that the high transfection efficiency of ONE came at the greatest cost to cell survival, and that MRNA preserves CAR T cell viability better than the other systems. In addition, prolonged transposase expression can cause cell stress and death, whereas the transient expression in MRNA reduces this stress and yields more viable cells.^16^

The TCN is an important parameter for the CAR T cell safety profile, since a high TCN is theoretically linked to insertional mutagenesis and is associated with the development of lymphoma from the edited cells following adoptive transfer^17^. The US FDA used to recommend that for therapeutic products with long-term integration, the integration copy number should not exceed 5 copies per cell.^18^ This was later amended to be looser, shifting to a release criterion that is based on product-specific risk stratification.^19^ Nevertheless, in its current absolute count, ONE showed a high transgene copy number, which is clinically inappropriate and carries an elevated risk of insertional mutagenesis^17^. In contrast, MRNA had the lowest copy number at all concentrations and should be the safest in terms of transgene integration.^12,16^.

Ideal point and Euclidean distance analysis suggest that ONE is the best format when a feeder is utilized, whereas MRNA is the best under the feeder-free condition. This result is consistent with that of King et al., who found that combining multiple genes into a single plasmid yields better efficiency than transfecting two or three separate plasmids.^20^ The reason proposed is that plasmids are taken up unequally by cells, so the higher the number of plasmids, the lower the likelihood that a cell receives all of them.

## Materials and Methods

### Plasmid construction

To construct a plasmid for mRNA transposase production, piggyBac transposase (pBase) was first amplified from the pCMV-piggyBac plasmid. Subsequently, both the amplified pBase and the pUC-ccTEV-CD117 IVT vector^21^ were digested with XhoI and KpnI restriction enzymes. The pBase fragment was then subcloned into the pUC-ccTEV vector.

To generate a helper-independent one-plasmid system, the QuikChange site-directed mutagenesis technique was first employed to replace the BglII restriction site with an AfeI site on the pIRII-CD19CAR(ΔCH2CH3) plasmid^14^. Subsequently, the EF-1α promoter was amplified from the pHIV-EGFP plasmid, adding AfeI and PspXI cut sites incorporated at the ends of the PCR product. The amplified EF-1α promoter was then subcloned into the QuikChange-modified pIRII-CD19CAR(ΔCH2CH3) plasmid.

### *In vitro* transcription (IVT)

The plasmid encoding pBase was digested overnight using XhoI and KpnI-HF (New England Biolabs, Ipswich, MA, USA). Following digestion, the DNA was precipitated with isopropanol. *In vitro* transcription was performed using the MEGAscript™ T7 Transcription Kit (Thermo Fisher Scientific, Waltham, MA, USA), CleanCap Reagent AG (3’ OMe), and N1-methylpseudouridine-5’-triphosphate (both from TriLink BioTechnologies, San Diego, CA, USA) in place of UTP, to generate 5’ capped mRNA encoding pBase. The transcription reaction was conducted for 3 hours at 37 °C with 5 µg of linearized plasmid template per 100 µL reaction. After transcription, double-stranded RNA (dsRNA) contaminants were removed using a cellulose-based purification method, described previously.^22^

### Peripheral Blood Mononuclear Cells (PBMCs) isolation and T cells selection

Peripheral blood mononuclear cells (PBMCs) were extracted from buffy coats of healthy donors by density gradient centrifugation with Ficoll-Paque Premium (Cytiva). All participants provided informed consent in line with ethical guidelines. T cells were enriched using CD4/CD8 Microbeads (Miltenyi Biotec, Bergisch Gladbach, Germany) with LS column and directly magnetic labeled. CD4+ T cells and CD8+ T cells were subsequently isolated.

### T cell activation

After PBMCs isolation and T cell selection, T cells were activated in a 24-well plate using T cell TransAct^TM^ (Miltenyi Biotec) and expanded in complete TexMACS medium (Miltenyi Biotec) supplemented with 5% FBS, 100 U/mL penicillin/streptomycin, 10 ng/mL human IL-7, and 5 ng/mL human IL-15 (both Miltenyi Biotec) and cultured under standard conditions (37 °C, 5% CO_2_, in a humidified incubator) for 3-4 days before electroporation.

### T cell electroporation

For electroporation, T cells were transfected with IVT mRNA pBase combination with one donor plasmid, helper-and-donor one-plasmid system, conventional two-plasmid system. T cells were then electroporated with 4D-Nucleofector X Unit and 20 µL Nucleocuvette Strips (both Lonza, Basel, Switzerland). Approximately 4.0 × 10^6^ of T cell were resuspended in 20 µL of P3 Primary Cell Solution (Lonza) containing plasmid DNA or a mixture of plasmid DNA with mRNA transposase and then electroporated using EO-115 pulse code. Then, the cells were transferred to complete cell media supplemented with 5% FBS, 100 U/mL penicillin/streptomycin, 10 ng/mL human IL-7, and 5 ng/mL human IL-15 (both Miltenyi Biotec) and cultured under standard conditions (37 °C, 5% CO_2_, in a humidified incubator).

### CAR T cells stimulation using feeder cells

The non-T cell population obtained from T cell selection was collected. Subsequently, 1.0 × 10^6^ cells were resuspended and irradiated with 25 Gy γ-irradiation. The irradiated non-T cells were used as feeder cells by mixing them with the electroporated T cell population and cultured under standard conditions.

### Flow cytometry analysis

To assess the transfection efficiency, the transfected cells were analyzed using MACSQuant flow cytometer (Miltenyi Biotec). CAR expression was detected using APC-anti-human CD3 Antibody (clone OKT3, #317318; Biolegend) in combination with PE-Labeled Monoclonal Anti-FMC63 scFv Antibody (#FM3-HPY53; ACROBiosystems, Newark, DE, USA). To assess cell viability, 7-Amino-Actinomycin D (7-AAD)(BD Biosciences) was used to select live cells.

To assess exhaustion markers, subset, and immunophenotype of CAR T cells, approximately 2.0 × 10^5^ cells were harvested and stained with PE-Cy^TM^7 Mouse Anti-Human CD279 (PD-1) (clone EH12.1, #561272; BD Biosciences), BB515 Mouse Anti-Human TIM-3 (CD366) (clone 7D3, #565558; BD Biosciences), BV480 Mouse Anti-Human TIGIT (clone TgMab-2, #570376; BD Biosciences), BV421 Mouse Anti-Human LAG-3 (clone T47-530, #565720; BD Biosciences), BB515 Mouse Anti-Human CD4 (clone RPA-T4, #564419; BD Biosciences), PE-Labeled CD8 (clone SK1, #340046; BD Biosciences), BV510 Mouse Anti-Human CD45RO (clone UCHL1, #563215; BD Biosciences), and BV421 Mouse Anti-Human CD62L (clone DREG-56, #563862; BD Biosciences).

For isotype controls, the following antibodies were used: BV421 Mouse IgG2b, k isotype Control (clone 27-35, #562748; BD Biosciences), BV510 Mouse IgG2a, k isotype Control (clone MOPC-173, #563483; BD Biosciences), PE-Cy^TM^7 Mouse IgG1, k isotype Control (clone MOPC-21, #565573; BD Biosciences), FITC Mouse IgG1, k isotype Control (clone MOPC-21, #400108; BD Biosciences).

### Transgene copy number detection using droplet digital PCR (ddPCR)

For transgene copy number analysis in transfected CAR-T cells, genomic DNA was extracted using a DNA Mini Kit (QIAGEN, Hilden, Germany). Concentrations lower than or equal to 60 ng/µL were considered adequate. Samples more concentrated than 60 ng/µL were diluted to a final concentration of less than 60 ng/µL.

Reactions were performed with the ddPCR supermix for probes (No dUTP), ddPCR CNV HEX-200R Assay (both BIO-RAD, Hercules, CA, USA), a primer pair, and the DNA template. To prepare droplets for the ddPCR reactions, a mixture was prepared and added with oil to the QX200 Droplet Generator (BIO-RAD). The plate containing these droplets was heat sealed using a PX1 PCR Plate Sealer (BIO-RAD) at 180°C for 5 sec and was loaded into a T100 Thermal64 Cycler (BIO-RAD). The sample was denatured for 10 min at 95°C and then subjected to 40 cycles of 30 sec at 94°C followed by 60 sec at 58.4°C. Finally, the droplets were cooled to 4°C to stabilize them. The plate was then analyzed in QX200 Droplet Reader (BIO-RAD). Positive and negative droplets were analyzed based on fluorescence amplitude difference. For each condition, technical triplicates were run, and the mean was used.

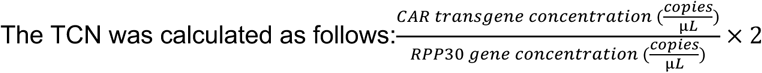

### Data analysis

Data analyses were performed on GraphPad Prism (version 8.4.0). The data are expressed as mean ± standard error of the mean (SEM). Group differences were assessed for statistical significance using two-way analyses of variance (ANOVA), with importance set at P values less than 0.05. *p < 0.05, **p < 0.01, ***p < 0.001, ****p < 0.0001; ns, not significant.

To identify optimal nucleic acids and delivery systems using ideal point and Euclidean distance, Z-score normalization was first performed on transfection efficiency, cell viability, and transgene copy number. The ideal point was defined as the maximum observed transfection efficiency and cell viability, but the minimum observed in transgene copy number. The optimal nucleic acid and delivery formats are those with the lowest Euclidean distances to the ideal point.

### Ethical consideration

This study, involving healthy human subjects, was approved by the Institutional Review Board of the Faculty of Medicine, Chulalongkorn University, COA no. 1424/2025. All participants were informed and consented to participate in the research. Such consent can be withheld or withdrawn by the participants at any time without justification. The study was conducted in accordance with the Declaration of Helsinki.

## Artificial intelligence disclosure

Claude Sonnet 5 was consulted for word choice in some sentences. No sentence was wholly generated by AI. The authors fact-checked all outputs from generative AI.

## Data availability statement

All data generated in this study are anonymized and will be available on Zenodo upon publication at https://doi.org/10.5281/zenodo.23010345

## Acknowledgments

This study was funded by the Thailand Science Research and Innovation Fund, Chulalongkorn University, grant no. HEA_FF_68_239_3000_038. The authors would like to thank Dr. Felipe Honjo Ide for helpful suggestions on multi-objective optimization. Figures 1B and 5A were created using BioRender.

## Author Contributions

TC conceptualized the project. YN supplied the original plasmids and the methods to utilize them. ND performed the experiments and formal analyses and wrote the original manuscript. TC, KS, and ST validated the results. TC reviewed and edited the manuscript. KS, ST, NH, and TC supervised the project and acquired funding.

---

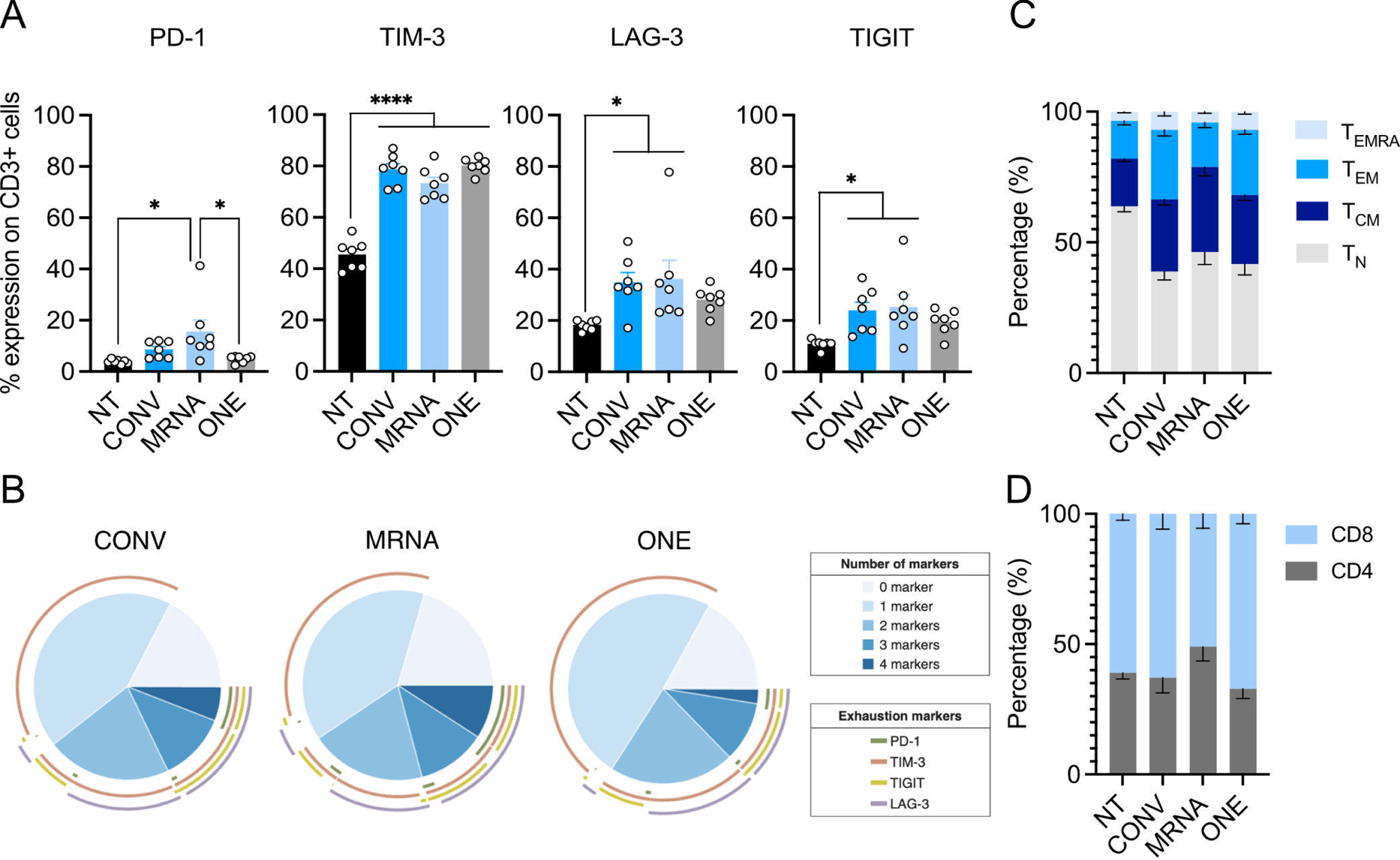

