## Supplemental Data for "PiggyBac transposon and transposase delivered in *trans*, in *cis*, and as mRNA for engineering CAR-T cells: a multi-objective optimization"

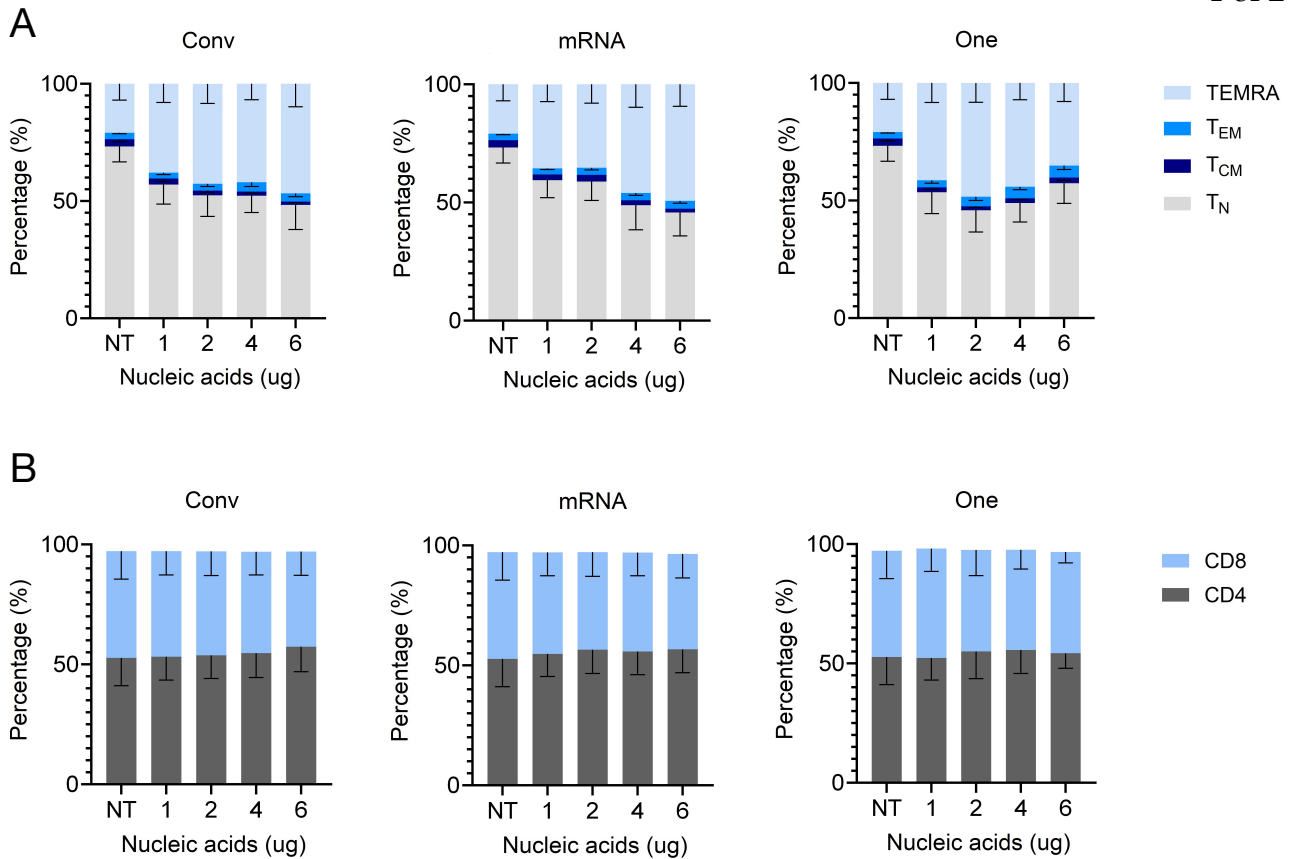

**Figure S1: Memory subsets and CD4/CD8 ratios of different deliveries under feeder-free condition.**

(A and B) Stacked bar chart showing memory subsets (A) and CD4/CD8 ratios (B)

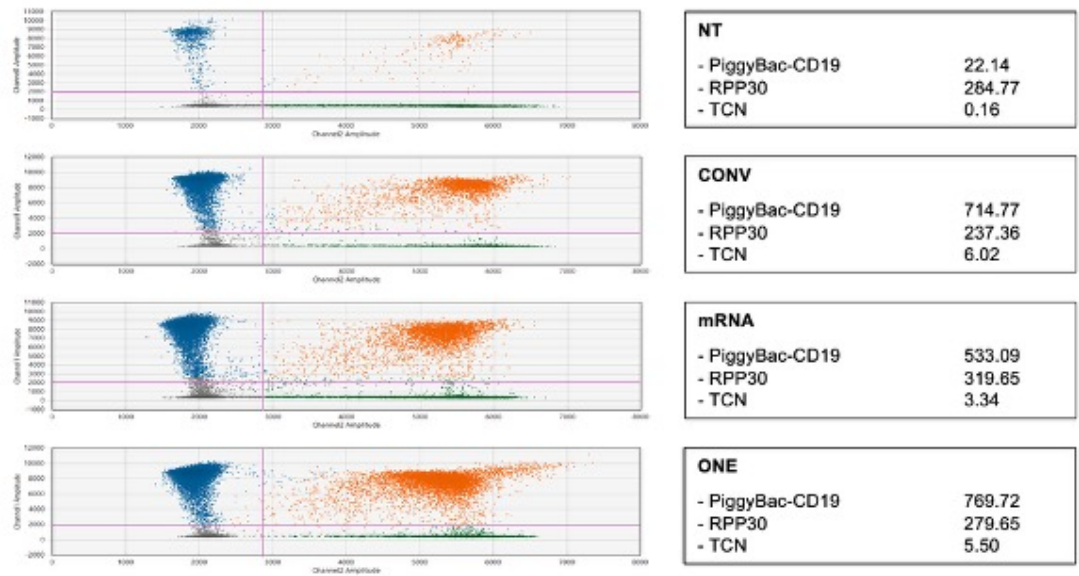

**Figure S2: Representative scatter plot from transgene copy number analysis of different deliveries with feeder enrichment.**

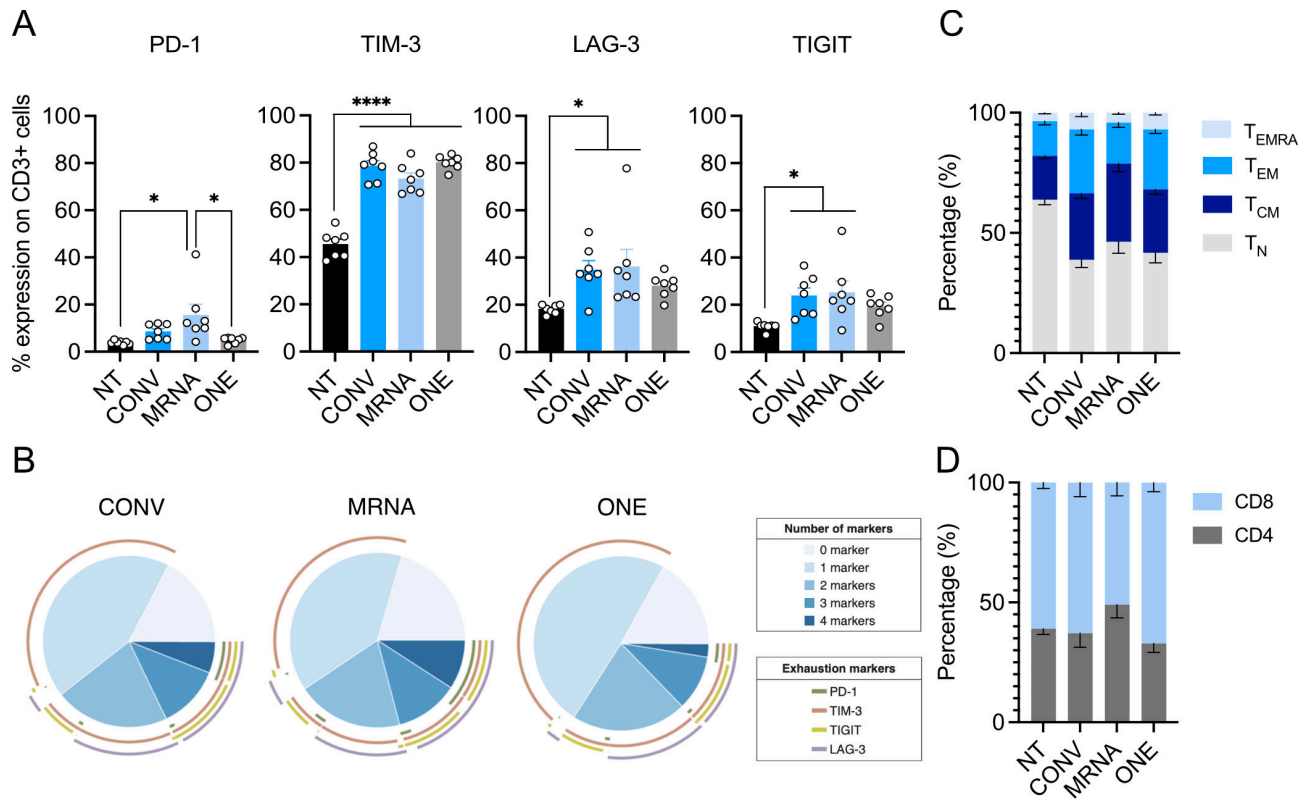

**Figure S3: Phenotypes of piggyBac-engineered CD19 CAR-T cells enriched with feeder cells.**

(A) CD19 CAR-T cell exhaustion marker expression, including PD-1, TIM-3, TIGIT, and LAG-3 at day 14 post-transfection. (B) Pie chart representing co-expressions of exhaustion markers. (C) Immunophenotype. (D) Subset. Data are presented as mean ± standard error of the mean (SEM), $n=7$. Data were analyzed by two-way ANOVA followed by Tukey's multiple-comparisons test. ns (not significant), \*$p<0.05$, \*\*$p<0.01$, \*\*\*$p<0.001$, \*\*\*\*$p<0.0001$.
